# Actin isovariant ACT7 controls root meristem development in Arabidopsis through modulating auxin and ethylene responses

**DOI:** 10.1101/2021.05.12.443766

**Authors:** Takahiro Numata, Kenji Sugita, Arifa Ahamed Rahman, Abidur Rahman

## Abstract

Meristem is the most functionally dynamic part in a plant body. The shaping of the meristem requires constant cell division and cell elongation, which are influenced by hormones and cell cytoskeletal component, actin. Although the roles of hormones in modulating meristem development have been extensively studied, the role of actin in this process is still elusive. Using the single and double mutants of the vegetative class actin, we demonstrate that actin isovariant ACT7 plays an important role in root meristem development. In absence of ACT7, but not ACT8 and ACT2, depolymerization of actin was observed. Consistently, *act7* mutant showed reduced cell division, cell elongation, and meristem length. Intracellular distribution and trafficking of auxin transport proteins in the actin mutants revealed that ACT7 specifically functions in root meristem to facilitate the trafficking of auxin efflux carriers PIN1 and PIN2, and consequently the transport of auxin. Compared with *act7*, *act7act8* double mutant shows slightly enhanced phenotypic response and altered intracellular trafficking. The altered distribution of auxin in *act7* and *act7act8* affects the response of the roots to ethylene but not to cytokinin. Collectively, our results suggest that ACT7 dependent auxin-ethylene response plays a key role in controlling Arabidopsis root meristem development.

## Introduction

Meristem, which sustains a reservoir of niche cells at its apex, is the most functionally dynamic part in a plant body. The formation of meristem is essential for plant growth and development, and this process is regulated by the interaction of multiple signaling networks, including receptor kinase signaling, transcriptional signaling, and hormonal signaling and transport (Barton, 2010). The shaping up of the meristem requires a constant cell division and cell elongation of the source stem cells. Both the hormones and cell cytoskeletal components have been attributed to regulate both cell division and cell elongation processes in the meristem cells (Rahman et al., 2007) Several studies have revealed that the asymmetric cell division and expansion processes are controlled by a dynamic cell cytoskeleton consisting of dozens of interacting proteins that responds to both environmental and developmental cues (Clayton and Lloyd, 1985; Lloyd, 1991; Baluška et al., 2001; Kandasamy et al., 2007, reviewed in Zhu and Giesler, 2015; Garcia-González and van Gelderen, 2021). Actin, a highly conserved and ubiquitous protein, is a major component of the cell cytoskeleton and essential for various important cellular processes, including cell division and cytokinesis, the establishment of cell polarity, cell division plane determination, cell shape, cell polarity, protein trafficking, cytoplasmic streaming, organelle movement and tip growth (Staiger and Schliwa, 1987; Smith and Oppenheimer, 2005; Staiger and Blanchoin, 2006; Kandasamy et al., 2009; Pollard and Cooper, 2009). Plants have multiple actin isoforms encoded by large ancient gene families (McLean et al., 1990; Meagher, 1991). In Arabidopsis, there are eight actin isovariants, which are grouped in two ancient classes, reproductive and vegetative classes (Kandasamy et al., 2009, ). Several lines of evidence strongly suggest that these two major classes of plant actin isovariants are functionally distinct. The vegetative actin isovariants, expressed in young tissues play roles in meristem development, while the plant’s reproduction is regulated by the reproductive class of actin (Gilliland et al., 2002). Previous experimental approaches using different isovariants of vegetative class actin further confirm that these three actin isovariants play distinct subclass-specific roles during plant morphogenesis (Gilliland et al., 2003; Kandasamy et al., 2009). For instance, among three vegetative actin, ACT7 had been shown to be preferentially and strongly expressed in all young and developing vegetative tissue including root meristem, and *ACT7* expression is strongly altered by plant hormone auxin and several external stimuli (McDowell et al., 1996a). It had also been shown that ACT7 regulates germination, root growth, epidermal cell specification, root architecture, and root thermomorphogenesis (Gilliland et al., 2003; Kandasamy et al., 2009, Parveen and Rahman, 2021). On the other hand, ACT2 and ACT8 regulate bulge site selection, root hair growth, and leaf morphology (Ringli et al., 2002; Nishimura et al., 2003; Kandasamy et al., 2009; Vaškebová et al., 2018).

Another major regulator of plant growth and development is hormones. The root meristem development is controlled by a complex regulatory network of hormones involving auxin, ethylene, and cytokinin (Bertell and Eliasson, 1992; Liu et al., 2017). Auxin plays a central role in this regulatory network through interacting with the other two hormones (Rahman, 2013). Polar auxin-driven formation of robust auxin gradient is an absolute requirement for root meristem development (Benková et al., 2003; Friml, 2003; Grieneisen et al., 2007). It had also been shown that root-produced auxin is required for root stem cell niche maintenance (Brumos et al., 2018a). Collectively, local auxin biosynthesis and auxin transport act redundantly to establish and maintain the robust auxin maxima critical for root meristem development. This auxin maxima at the root tip, in combination with separable roles of auxin in cell division and cell expansion, can explain the root meristem development.

Cytokinin is another major regulator to control the root meristem activity. Experiments resulting in depletion of cytokinin in the root meristem along with cytokinin mutants analyses revealed that cytokinin functions at the transition zone to control the cell differentiation rate (Dello Ioio et al., 2007; Müller and Sheen, 2007; Dello Ioio et al., 2008). The classical antagonistic interaction between cytokinin and auxin that regulates the root and shoot organogenesis has been extended to the root meristem development and shown to be regulated through a simple regulatory circuit converging on the *Aux/IAA* gene *SHY2/IAA3*, and polar auxin transport mediated auxin gradient (Dello Ioio et al., 2008a; Chapman and Estelle, 2009; Růzǐčka et al., 2009; Moubayidin et al., 2010a) Ethylene also plays an important role in root meristem development. Cell division in quiescent cells, which act as a source of stem cells, has been shown to be modulated by ethylene. The ethylene-induced new cells in the quiescent center (QC) express QC-specific genes that can repress the differentiation of surrounding cells (Ortega-Martínez et al., 2007). Consistently, constitutive triple response mutant *ctr1* shows reduced meristematic cell division and meristem size, while the ethylene insensitive mutant *ein2* shows the opposite phenotype. Exogenous application of ethylene also mimicked the constitutive phenotype (Street et al., 2015). Auxin-ethylene interaction for root growth and development has been well studied (Rahman et al., 2001; Rahman et al., 2002; Stepanova et al., 2005; Muday et al., 2012). Reduction in intracellular auxin level results in ethylene insensitivity that affects the root morphology. The ethylene sensitivity and the root morphology could be restored by increasing the intracellular auxin level using exogenous auxin (Rahman et al., 2001). Conversely, ethylene has been shown to promote auxin synthesis and transport in root tips resulting in the formation of a second auxin maximum in the root elongation zone and alteration in root meristem development (Růzicka et al., 2007; Stepanova et al., 2007; Swarup et al., 2007; Stepanova et al., 2008). Ethylene-induced local auxin synthesis is achieved by upstream tryptophan biosynthetic genes *WEI2/ ASA1 and WEI7/ ASB1* that are precisely regulated by ethylene (Stepanova et al., 2005; Okamoto et al., 2008). This local auxin synthesis by ethylene also alters the auxin gradient in the root tip and affects the root meristem development (Okamoto et al., 2008). Furthermore, it has been shown that *SHY2/IAA3* is a point of convergence for both ethylene and cytokinin that negatively regulates cell proliferation (Street et al., 2015).

Although the cell division and cell proliferation that control the meristem development are influenced by both hormones and cell cytoskeleton component actin, the integrating mechanisms of these two components remain elusive. It has been shown that mutations in vegetative actin isovariant genes result in alteration in root morphology and root development (Ringli et al., 2002; Gilliland et al., 2003; Nishimura et al., 2003; Kandasamy et al., 2009; Vaškebová et al., 2018). However, it remains obscure how the isovariant specific actin affects the root developmental process. In the present study, using a combinatorial approach of physiology, genetics, and cell biology, we demonstrate that actin isovariant ACT7 plays an important role in maintaining root meristem development. In absence of ACT7, but not ACT8 and ACT2, depolymerization of actin was observed. Consistently, *act7* mutant showed reduced cell division, cell elongation, and meristem length. Intracellular distribution and trafficking of auxin transport proteins in the actin mutants revealed that ACT7 specifically functions in root meristem to facilitate the trafficking of auxin efflux carriers PIN1 and PIN2, and consequently the transport of auxin. Compared with *act7*, *act7act8* double mutant shows slightly enhanced phenotypic response and altered intracellular trafficking. The altered distribution of auxin affects the root’s response to ethylene but not to cytokinin. Collectively, our results suggest that Arabidopsis root meristem development is affected by ACT7 isovariant-mediated modulation of auxin-ethylene responses.

## Results

### Root actin organization is primarily controlled by actin isovariant ACT7

To investigate the effect of mutations in vegetative actin genes on cellular actin structure and organization, we observed the actin structure using both immunostaining and live-cell imaging approaches. Actin in elongation zone was observed as the structure of the actin in this zone is most reliable to compare. In this study, we used the alleles of the actin mutants, namely *act2-1, act7-4 and act8-2, and act7-4act8-2*, which were extensively characterized at physiological, molecular, and protein levels (McKinney et al., 1995; McDowell et al., 1996a; Kandasamy et al., 2009). For live-cell imaging, actin mutants were crossed with the actin marker line ABD2-GFP (Actin Binding Domain 2 of Fimbrin protein, Wang et al., 2007) and homozygous lines were selected. The actin structure observed by both approaches was largely similar. In wild type, actin labeled in elongation zone showed fine filamentous cable-like structure (Fig. 1A). Actin filament structure in *act2-1 and act8-2* showed normal actin filaments like wild-type (Figs. 1B, 1D). Comparable results were observed with live-cell imaging using actin marker lines (Figs. 1G, 1H, 1J). Loss of both ACT2 and ACT8 did not affect the structure of the actin. Although a slight reduction of actin was observed in immunostained samples, the actin structure in *act2act8*ABD2-GFP lines was like wild-type (Figs. 1E and 1K). In contrast, the loss of ACT7 drastically affected actin structure, showing segmented and aberrant actin cables (Figs. 1C, 1I). This fragmented and aberrant actin phenotype was slightly more enhanced in *act7act8* double mutant (Figs. 1F; 1L). To further confirm the implications of the mutations in actin structure, we quantified four key parameters that are used for actin quantification namely: (1) occupancy, representing the filament density, (2) skewness, representing the filament bundling, (3) Δθ, representing average angle against the longitudinal axis, and (4) normAvgRad, representing the parallelness of the filamentous actin (Higaki et al., 2010; Ueda et al., 2010). Occupancy was found to be altered in *act7* and *act2* mutants. *act2* showed a slightly higher occupancy compared with wild-type, while *act7* showed a slightly lower occupancy (Fig. 1M). No difference was observed for skewness across the mutant lines (Fig. 1N). Filament orientation and filament paralleleness were found to be altered in *act7* and *act7act8* compared with wild-type and other actin mutants (Figs. 1O-P), which is consistent with the observed segmented and aberrant actin cables in these two mutants by immunostaining and live-cell imaging. Collectively, these results suggest that ACT7 plays a key role in maintaining the structural integrity of root actin.

**Figure 1.**
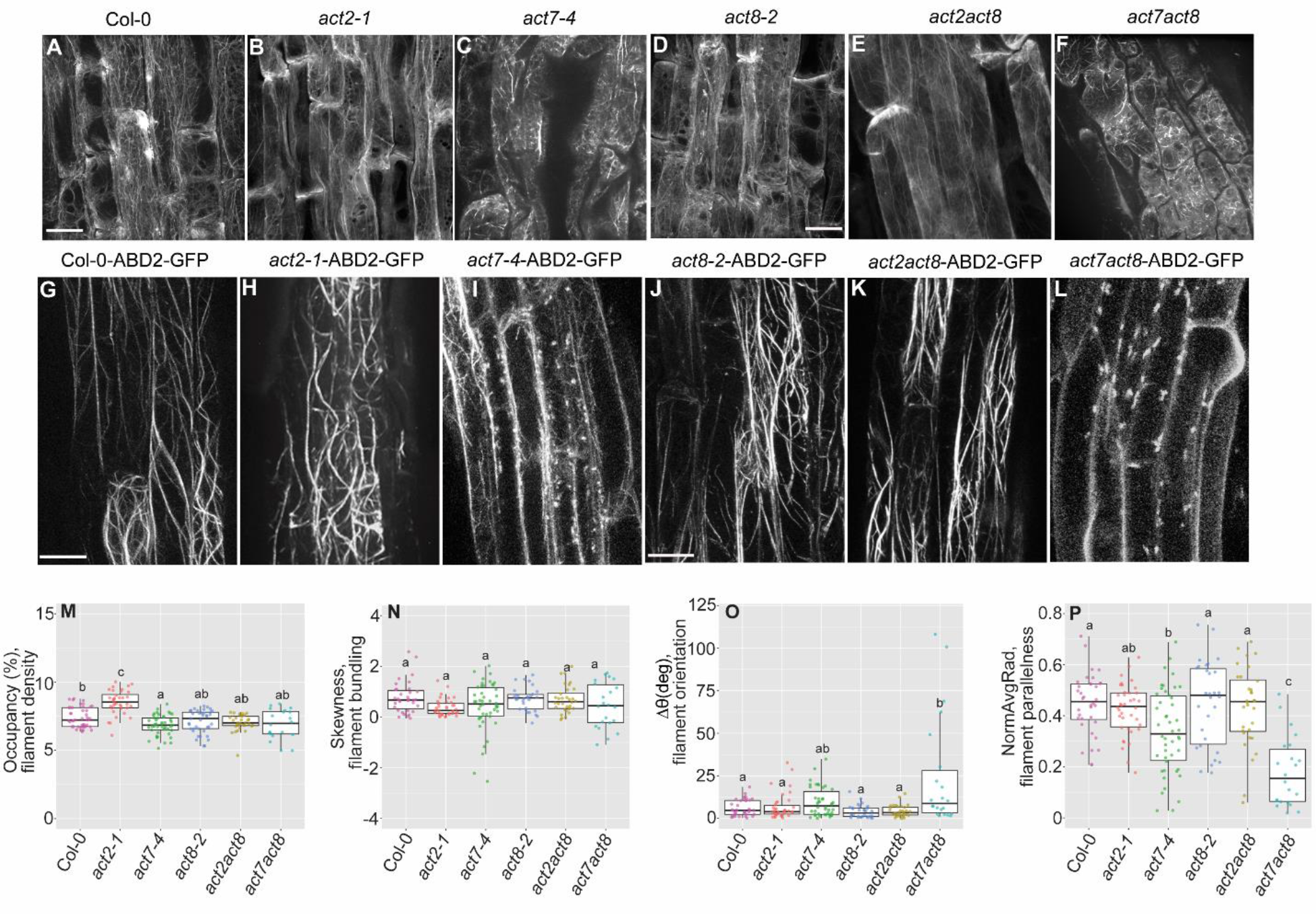
Effect of loss of Actin isovariant proteins on intracellular actin organization. Upper panel (A-F) chemical fixation. Lower panel (G-L) live-cell imaging. The images are representative of at least three fixation runs, with 5–7 roots per genotype in each run. Five-day-old Arabidopsis roots were fixed (for chemical fixation) or mounted in liquid growth medium on a cover glass (for live-cell imaging) and actin was localized using confocal laser microscopy. The images were obtained from the elongation zone of the root and imaged using the same confocal settings for immunostaining and live-cell imaging, respectively. Images are projections of 10–12 optical sections. Bars=10 μm for upper panel and 20 μm for lower panel. Quantification of actin filaments (M-P) (M) percent occupancy, (N) skewness, (O) NormAvgRad (P) Δθ. Cells obtained from live cell imaging experiment (G-L) were used for actin quantification. Vertical bars represent mean ± SE of the experimental means from at least three independent experiments (n = 3 or more), where experimental means were obtained from 20–30 cells. Comparisons between multiple groups were performed by analysis of variance (ANOVA) followed by the Tukey–Kramer test. The same letter indicates that there are no significant differences (P < 0.05).

### Cellular actin organization affects the primary root elongation and meristem development

To understand whether the change in the actin structure observed in actin mutants correlates with the plant growth and development, next we performed the comparative analysis of these actin isovariant mutants against wild-type for root elongation, meristem size, cell production rate, and cell length (Fig. 2, Table 1). The seedling phenotype of single and double mutants revealed that the primary root elongation is severely compromised in the mutants where ACT7 is mutated. Both the *act7* and *act7act8* mutants showed dwarf root phenotype, while the wild-type-like or a slight increase in the primary root elongation was observed in *act2*, *act8,* and *act2act8* (Fig. 2A). The *act7act8* double mutant showed a little more severe phenotype than the *act7-4* single mutant (Fig. 2A). These phenotypic observations suggest that the intracellular actin organization directly influences the primary root elongation and further reveal that that ACT7 is the primary modulator of this process.

**Figure 2.**
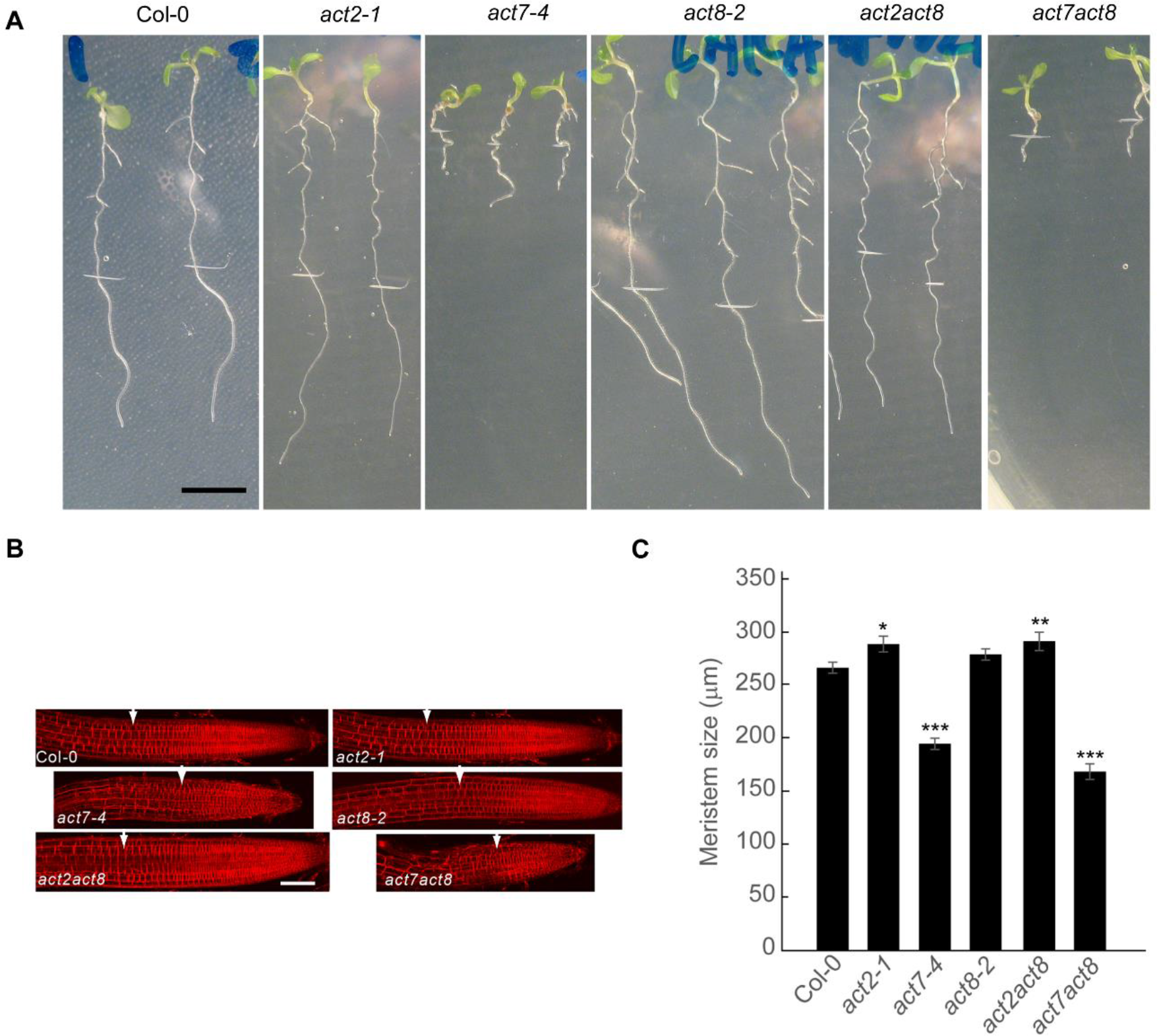
ACT7 plays an important role in determining primary root development and meristem size. (A) Seven-day-old seedling phenotype of actin single and double mutants. Loss of ACT7 drastically affects the seedling phenotype, including primary root growth. (B-C) Image and the size of the primary root meristem in actin isovariant mutants. For root meristem imaging, seven- day-old seedlings were stained in 2 μM FM 4-64 for 15 minutes and subjected to imaging under confocal laser microscope using the same settings. Images and data are representative of at least 3-4 biological replicates. Vertical bars represent mean ± SE of the experimental means obtained from at least 3 experiments (n= 3 or more), where experimental means were obtained from 6-8 seedlings per treatment. Asterisks represent the statistical significance between wild-type and mutants as judged by the Student’s *t*-test (*P<0.5, **P<0.01, ***P< 0.001). Bars, 5 mm (A); 50 μm (B). Arrowheads indicate the start of the transition zone.

**Table 1:** Actin isovariant ACT7 is the primary modulator of root elongation, cell length, and cell production in Arabidopsis

| Genotype | Primary root elongation (mm) | Cell length (μm) | Cell production rate |
| --- | --- | --- | --- |
| Col-0 | 9.05±0.03 | 166±1.5 | 54.0±2.2 |
| *act2-1* | 9.90±0.21 | 157±0.29* | 62.0±0.3* |
| *act7-4* | 3.39±0.21*** | 87±5.34*** | 39.5±2.2*** |
| *act8-2* | 8.90±0.16 | 166±6.8 | 52.4±1.3 |
| *act2act8* | 9.09±0.60 | 159±2.08 | 61.5±2.3 |
| *act7act8* | 1.8±0.02*** | 68±7.23*** | 26.8±2.2*** |

Data are means ± S.E of three replicate experiments. 4-day-old vertically grown seedlings were transferred to new plates and grown for another three days. The measurements reflect the behavior over the third day of treatment. Asterisks represent the statistical significance between wild-type and mutants as judged by the Student's *t*-test (\**P*<0.5, \*\*\**P*< 0.001).

Root elongation is regulated by both cell division and cell elongation and consequently linked to the root meristem size and development (Beemster and Baskin, 1998). The assessment of meristem length in vegetative class actin isovariant mutants revealed that the dwarf root phenotypes observed in *act7* and *act7act8* mutants are a consequence of reduced meristem development (Figs. 2B, 2C). In contrast, *act2* and *act2act8* showed a slight but statistically significant increase in meristem length (Figs. 2B, 2C) To further understand how actin isovariant regulates the root meristem size, we took a kinematic approach as described earlier (Rahman et al., 2007). The root elongation rate and the length of the newly matured cortical cells were measured in wild type, *act2-1*, *act8-2*, *act7-4*, *act2act8*, and *act7act8* mutants. The ratio of mature cell length to root elongation rate gives the time required to produce one cortical cell (per cell file); which is also defined as the cell production rate (Silk et al., 1989). The cell production rate represents the output of the meristem reflecting both the number of dividing cells and their rates of division (Beemster and Baskin, 1998; Rahman et al., 2007). Kinematic analysis revealed that the loss of ACT7 results in a significant reduction in root elongation, cell length, and subsequently lower cell production rate (Table 1). Further, the double mutant *act7act8*, which shows a little more severe root phenotype than *act7-4*, also shows a reduced cell division activity compared with *act7-4*. These data suggest that ACT7 plays a dominant role in controlling the cell division activity at the primary root meristem. Loss of ACT8 alone does not change the cell production activity but the loss of ACT8 in conjunction with ACT7 affects the roots cell division ability negatively, consistent with the idea that actin plays a significant role in controlling the cell division and ACT7 isovariant is the primary modulator of this process (Table 1). On the other hand, loss of ACT2 results in an increased cell production rate although the cell length is not affected. The double mutant *act2act8* also shows a similar trend (Table 1). The increased cell production rate in *act2* and *act2act8* mutants explains why these mutants show increased root meristem size compared with wild-type (Figs. 2B, 2C) Taken together, these results suggest that the root meristem development is controlled by isovariant specific actin, and ACT7 is the primary modulator of this process.

### ACT7 modulates the intracellular trafficking of a subset of auxin transport proteins

In eukaryotic cells, movement of the proteins to specific organelles or to and from the membrane is regulated by protein trafficking, an important process for cellular activity and growth. It is generally assumed that the cytoskeleton component actin acts as a network of tracks for the movement of vesicles between cellular compartments (Simon and Pon, 1996; Huang et al., 1999; Nebenführ and Staehelin, 2001; Kim et al., 2005). Polar transport and the formation of auxin gradient in the root meristem largely rely on the intracellular trafficking of PIN proteins through which the proteins are correctly localized to the membrane (Feraru and Friml, 2008), and this localization of PIN efflux carriers is shown to be actin-dependent. Disruption in actin structure results in reduced trafficking of these proteins to and from the membrane and subsequently affects their polar localization (Geldner et al., 2001; Muday and Murphy, 2002; Rahman et al., 2007; Kleine-Vehn and Friml, 2008). Since polar auxin transport mediated auxin gradient at root tip plays a major role in regulating the root meristem development, and actin mutants show altered root meristem size, we hypothesized that the observed meristem phenotype in these mutants may be linked to the altered trafficking of the auxin transporters. To test this hypothesis, we investigated the intracellular localization of the PIN proteins in actin mutants. Live-cell imaging analyses using translationally fused GFP with PIN proteins in various actin mutant backgrounds revealed that the loss of actin does not alter the polar localization of these proteins (Figs. 3-5). However, the intracellular trafficking of PIN1 and PIN2 was found to be affected by a specific actin isovariant ACT7. Intracellular PIN1 and PIN2 protein agglomerations were observed in *act7* and *act7act8* mutants but not in *act2*, *act8* or *act2act8* mutants (Figs. 3 and 4). The intracellular agglomeration of PIN1 and PIN2 resulted in reduced accumulation of PIN1 and PIN2 in the membrane. Compared with PIN2, the effect of loss of ACT7 was found to be more dominant in PIN1. Further, the PIN2 protein aggregation was observed only in the meristem cells of *act7* and *act7act8*, not in the cells in the transition zone (Fig. 4). For PIN7, loss of ACT7 results in a reduction in the expression of the protein, which is further downregulated in the *act7act8* double mutant. No notable change in PIN7 expression was observed in *act8* or *act2act8* mutant background (Fig. 5A). On the other hand, loss of ACT7 or ACT8 did not affect the trafficking or expression of PIN4 or auxin influx carrier AUX1 (Fig. 5B). The observed intracellular agglomeration in only GFP fused PIN1 and PIN2 lines but not in other GFP fused lines confirm that ACT7 specifically regulates the intracellular trafficking of these two proteins. To further confirm these results, cellular localization of PIN2 in *act* mutants was performed using anti-PIN2 antibody. As observed in the PIN2-GFP line, cellular agglomeration of PIN2 was observed in *act7* and *act7act8* mutants but not in *act8* mutant (Supplementary Figure 1).

**Figure 3.**
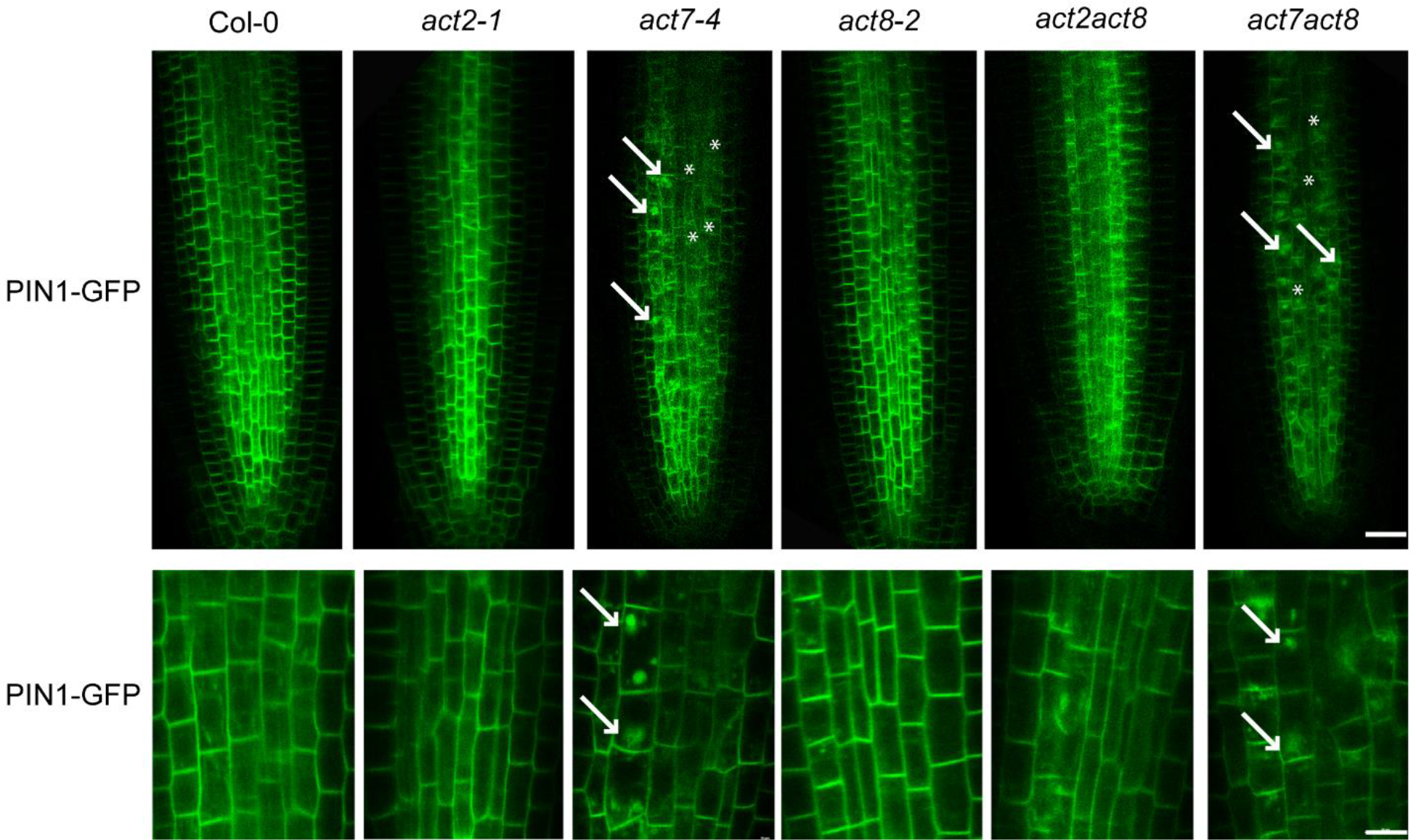
Loss of ACT7 but not ACT2 and ACT8 affects the intracellular dynamic cycling of PIN1. Seven-day-old PIN1:PIN1-GFP transgenic seedlings in wild-type and actin single and double mutant backgrounds were subjected to confocal imaging. The images were captured using the same confocal settings and are representative of 20 roots from three-four independent experiments. The lower panel represents the zoomed images. The intracellular agglomeration of PIN1, indicated by arrowheads, was observed exclusively in *act7* and *act7act8* mutants. The reduced signal in the membrane is indicated by the asterisks. Bars represent 10 μm.

**Figure 4.**
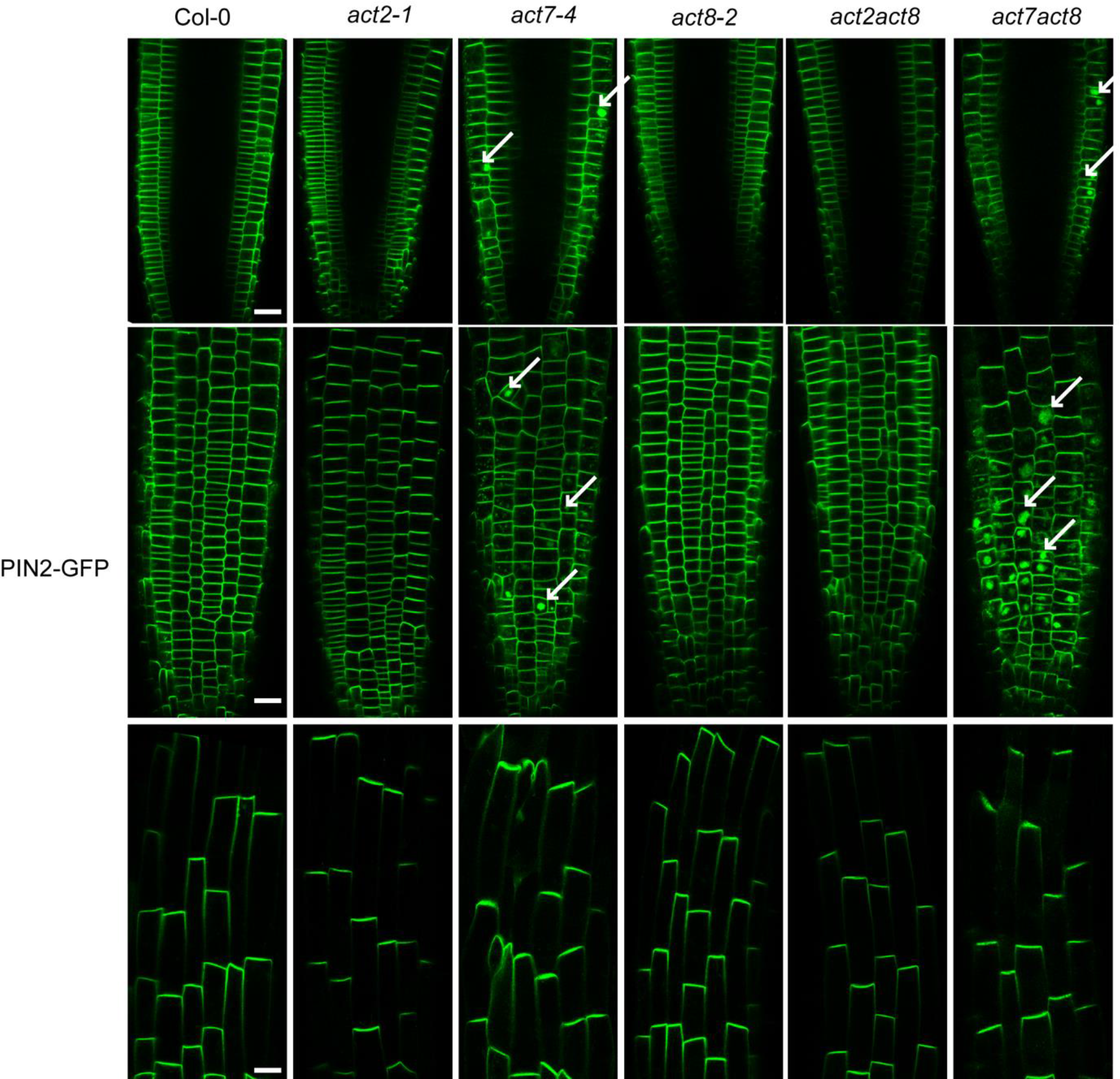
Loss of ACT7 but not ACT2 and ACT8 affects the intracellular dynamic cycling of PIN2. Seven-day-old PIN2:PIN2-GFP transgenic seedlings in wild-type and actin single and double mutant backgrounds were subjected to confocal imaging. The images were captured using the same confocal settings and are representative of 25 roots from three-four independent experiments. The upper and middle panels represent the images from the root meristem region. The lower panel represents the images from the transition zone. The intracellular agglomeration of PIN2 was observed exclusively in the meristematic region of *act7* and *act7act8* mutants but not in the transition zone (Bottom Panel). Bars represent 10 μm.

**Figure 5.**
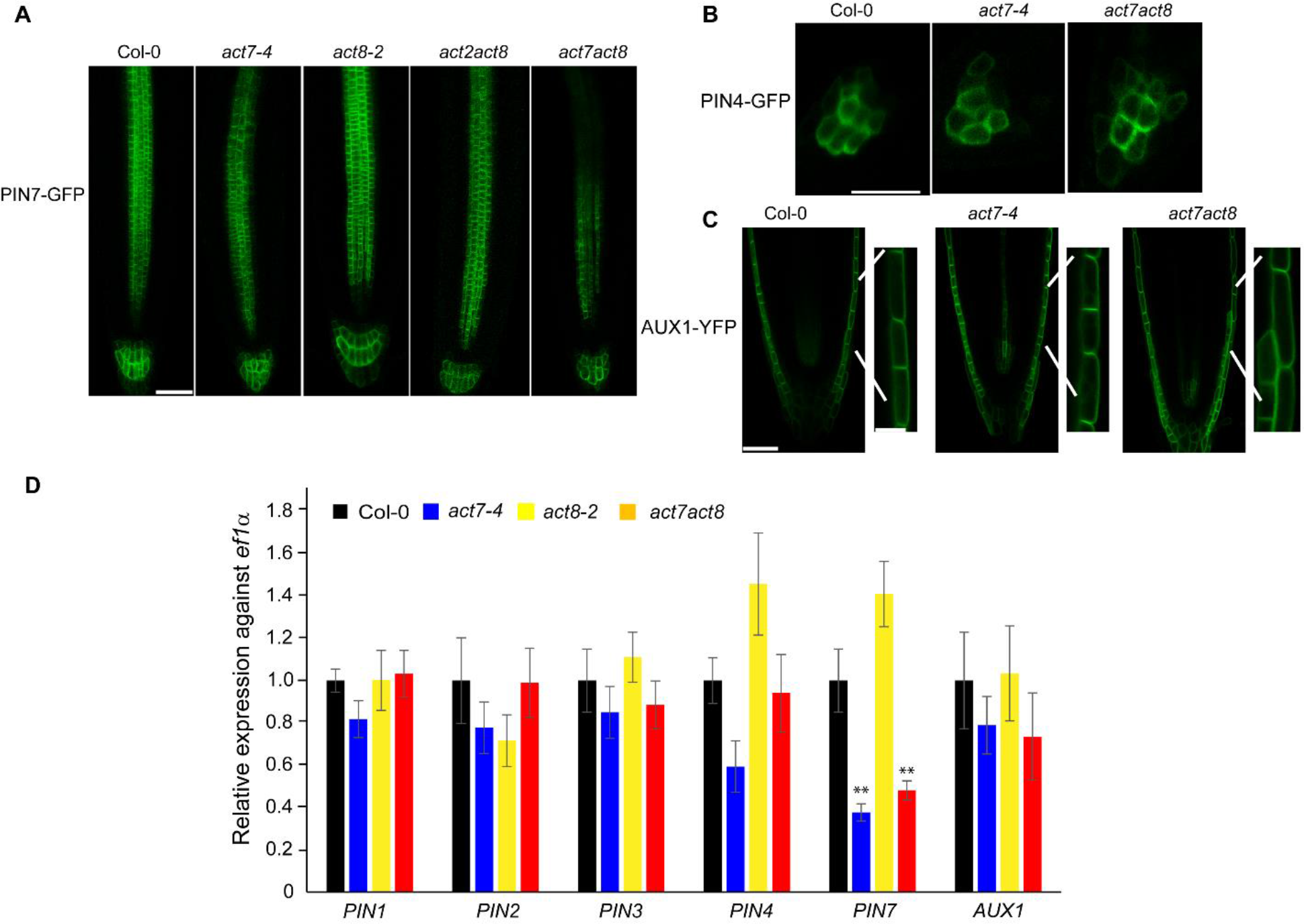
Effect of loss of ACT7 on PIN4, PIN7, and AUX1 expression Seven-day-old PIN7:PIN7-GFP, PIN4:PIN4-GFP, and AUX1-YFP transgenic seedlings in wild- type and actin single and double mutant background (A-C) were subjected to confocal imaging. The images were captured using the same confocal settings and are representative of 20 roots from three-four independent experiments. PIN7 expression was reduced in *act7* and *act7act8* mutants. PIN4 and AUX1 expressions were unaltered in *act7* and *act7act8* mutants. Bars represent 50 μm, and 10 μm for zoomed images of AUX1. (D) Relative quantification of *PINs* and *AUX1* in wild-type and actin single and double mutant backgrounds. Root samples were collected from seven-day-old light-grown seedlings for RNA isolation, cDNA preparation, and qRT-PCR analysis. All the data were normalized against *EF1α.* The data were obtained from at least three biological replicates (n=3 or more). Vertical bars in the graph represent mean ± SE. Asterisks represent the statistical significance between the means of wild type and mutants as judged by the Student’s *t-*test ( **P < 0.01).

The transcript analysis of *PINs/AUX1* further confirmed the observed changes at the protein level. Only *PIN7* expression was found to be downregulated in *act7* and *act7act8* mutants (Fig. 5C), while the expression of other *PIN*s and *AUX1* was unaltered by the loss of actin isovariants. Taken together, these results suggest that actin isovariant ACT7 possibly provides the primary track for the intracellular trafficking of PIN1 and PIN2.

### Auxin gradient at root tip is altered in *act7*and *act7act8* mutants

Since the intracellular trafficking of the auxin proteins play significant role in determining the cellular auxin flow, and the formation of auxin gradient is required for maintenance of stem cell niche and meristem development, we next investigated whether the observed alterations of intracellular trafficking of PIN1 and PIN2 affect the auxin gradient formation in actin isovariant mutants. Auxin gradient formation assessment using two widely used auxin reporter lines *DR5- GUS* and *IAA2-GUS* (Ulmasov et al., 1997; Luschnig et al., 1998) revealed that the actin isovariant ACT7 plays a major role in modulating this process as reduced GUS signal was observed at the *act7* root tip (Fig. 6). Consistently, auxin gradient formation was not altered in the *act8* mutant. *act7act8* double mutant showed a complete loss of auxin gradient at the root tip, although the GUS signal could be observed in the upper part of the root (Fig. 6). Exogenous application of IAA could increase the GUS signal at the root tip of *act7* but not in *act7act8* (Fig. 6). These results confirm that the root auxin gradient formation is primarily dependent on actin isovariant ACT7 modulated PIN1 and PIN2-mediated auxin flow.

**Figure 6.**
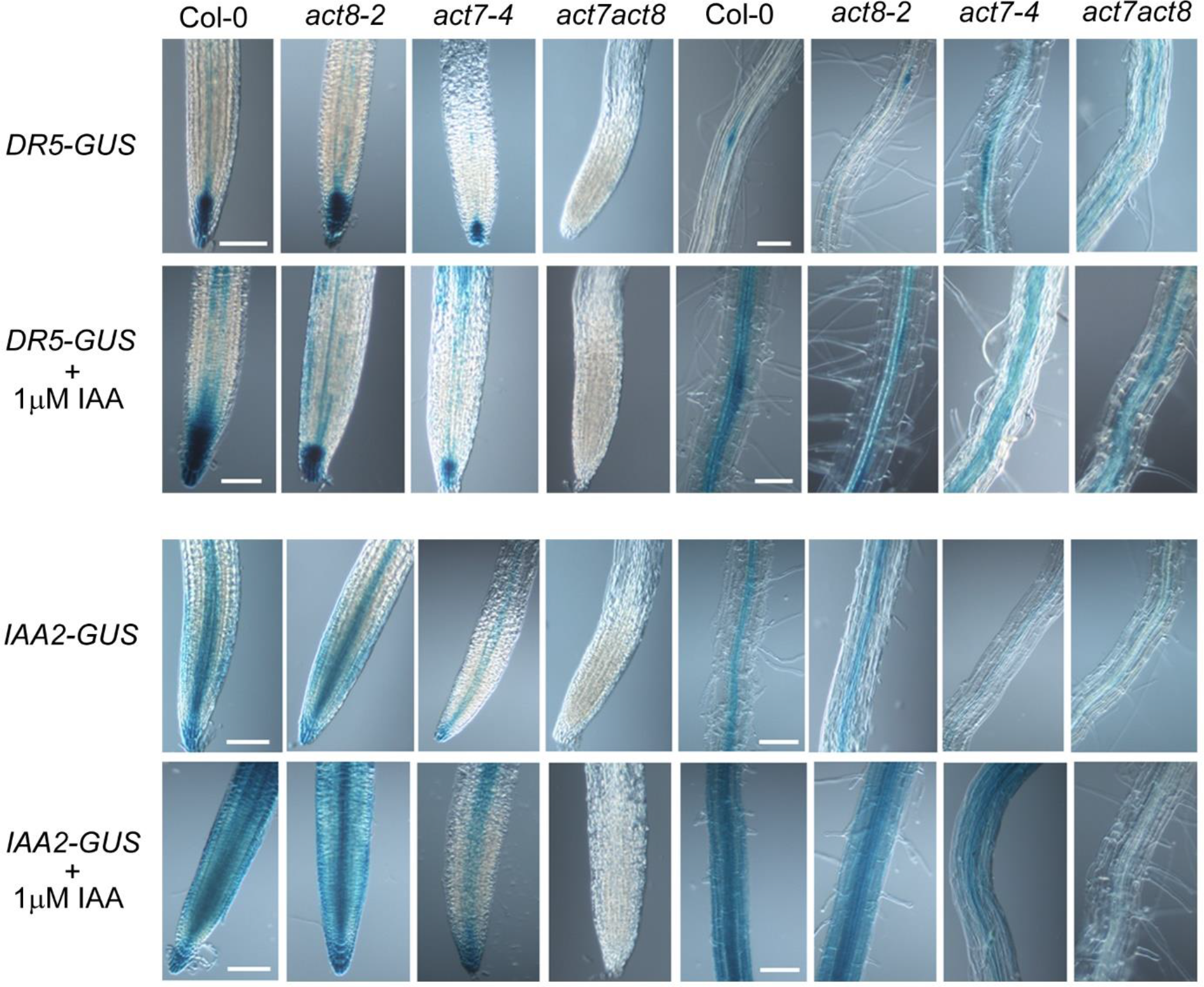
Effect of loss of actin isovariant on the root auxin gradient formation Five-day-old *DR5-GUS* and *IAA2-GUS* transgenic seedlings in wild-type and actin single and double mutant backgrounds were subjected to GUS staining. *DR5-GUS* transgenic seedlings were stained in buffer containing 1 mM X- gluc for 3 h and *IAA2-GUS* were stained for 1 h at 37℃. After the incubation, the roots were cleared for photography. Loss of ACT7 results in the formation of reduced auxin gradient at the root tip. Loss of both ACT7 and ACT8 further enhances the reduction. For exogenous auxin treatment, the seedlings were incubated in 1μM IAA for 3h and subjected to GUS staining and cell clearing. The images are representative of 20 seedlings from three-four independent experiments. The results were confirmed with two separate lines for each crossing. Bars represent 100 μm.

### Auxin transport is altered in *act7* mutant

To confirm that the reduced auxin gradient in *act7* mutant is due to the reduced transport of auxin resulting from the altered trafficking of PIN1 and PIN2, we performed the auxin transport assay using radiolabeled IAA. Compared with wild-type, both rootward and shootward transports were found to be reduced in *act7-4* mutant, confirming that the observed reduction in auxin gradient at root tip is due to the altered transport of auxin (Fig. 7).

**Figure 7.**
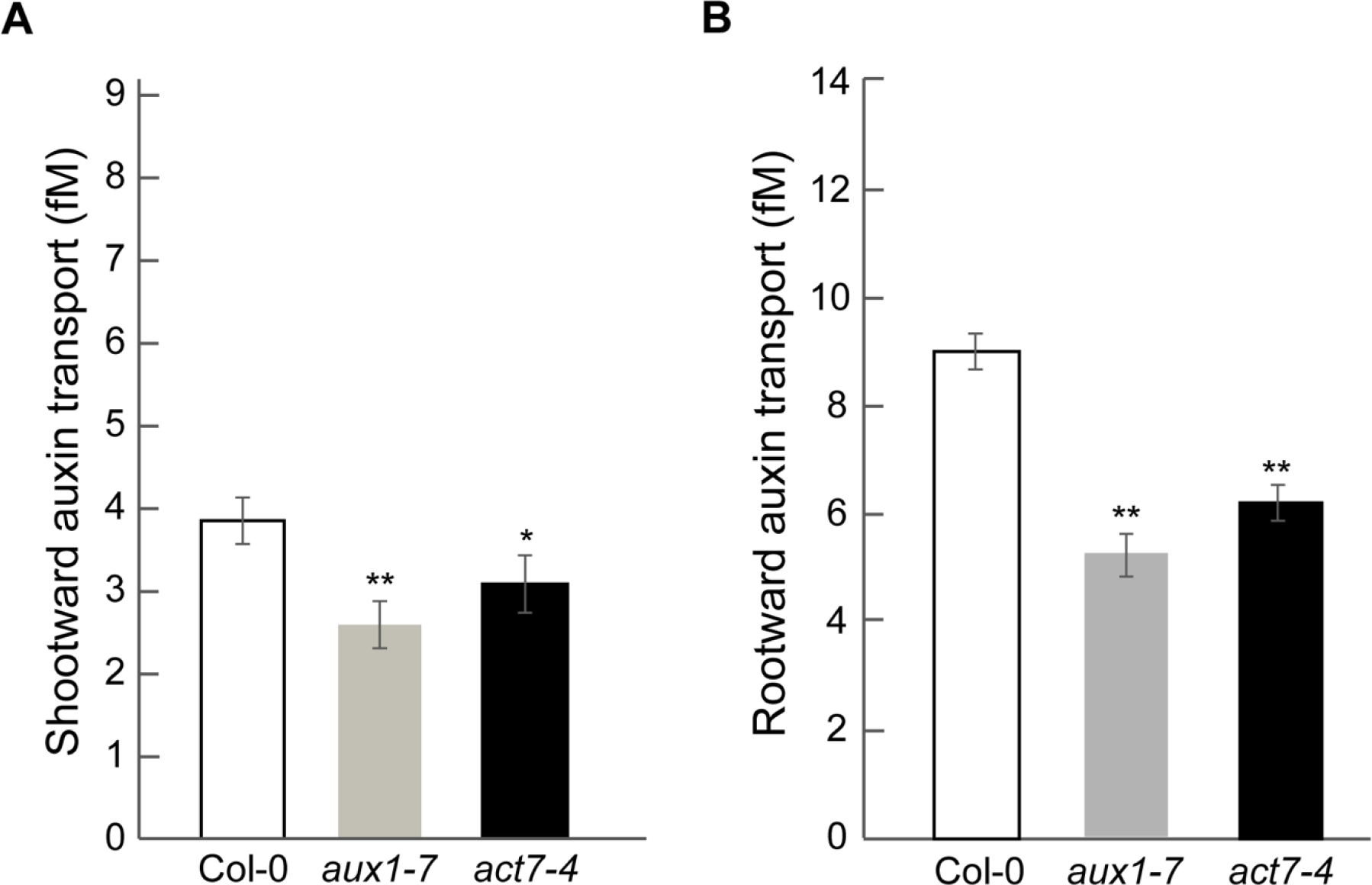
Auxin transport is disrupted in *act7* mutant. Shootward and rootward auxin transport were measured using five-day-old seedlings. The experiments were performed in triplicate and repeated at least three times. Auxin influx facilitator mutant *aux1-7* was used as a positive control. Asterisks represent the statistical significance between wild-type and mutants as judged by the Student’s *t*-test (*P<0.5, **P<0.01).

### Ethylene response is altered in *act7* and *act7act8* mutants

Ethylene can influence the auxin response by altering the local biosynthesis of auxin at the root tip (Stepanova et al., 2005; Stepanova et al., 2007; Swarup et al., 2007). Since ethylene is involved in regulating stem cell niche (Ortega-Martínez et al., 2007) and directly influences the root meristem cell division, we checked whether the reduced meristematic cell division observed in *act7* and *act7act8* mutants are also linked to root ethylene response. To clarify the issue, we investigated the expression of a reporter line containing a transcriptional fusion of *ANTHRANILATE SYNTHASE β1* to *GUS* (*ASB1-GUS*), which catalyzes one of the rate-limiting steps of tryptophan biosynthesis, a precursor of IAA (Stepanova et al., 2005), and is specifically induced by ethylene (Guo and Ecker, 2004; Stepanova et al., 2005; Okamoto et al., 2008). Consistent with the previous report, in the wild type, the GUS activity driven by *ASB1* was found to be exclusively localized at the root tip, and no expression in the elongation zone (Stepanova et al., 2005; Figure 8A). Loss of ACT7 and both ACT7 and ACT8 result in ectopic GUS expression. In *act7* and *act7act8* mutants, although a complete loss of GUS activity was observed at the root tip, unusual high activity was observed in the elongation zone, suggesting that ethylene-induced auxin production at the root tip is blocked in these mutant backgrounds (Fig. 8A). These results also further suggest that *ASB1* regulated tryptophan pool may serve as the sole source for the auxin produced at the root tip. To further understand whether the reduction in auxin response also affects the ethylene response, we investigated the expression of the *ETR2, CTR1,* and *EIN2*, that regulate the ethylene response pathway. Although *ETR2* and *CTR1* expressions were unaltered in *act7* and *act7act8* mutants, *EIN2*, which functions as a positive regulator for ethylene response (Alonso, 1999) was found to be down-regulated in *act7act8* mutant (Fig. 8B-D). To further understand the biological significance of these changes, we performed the root growth assay in presence of ACC, a precursor of ethylene. Although *act7* showed a wild-type like response to ACC-induced root elongation inhibition, *act7act8* showed a clear resistance (Fig. 8E).

**Figure 8.**
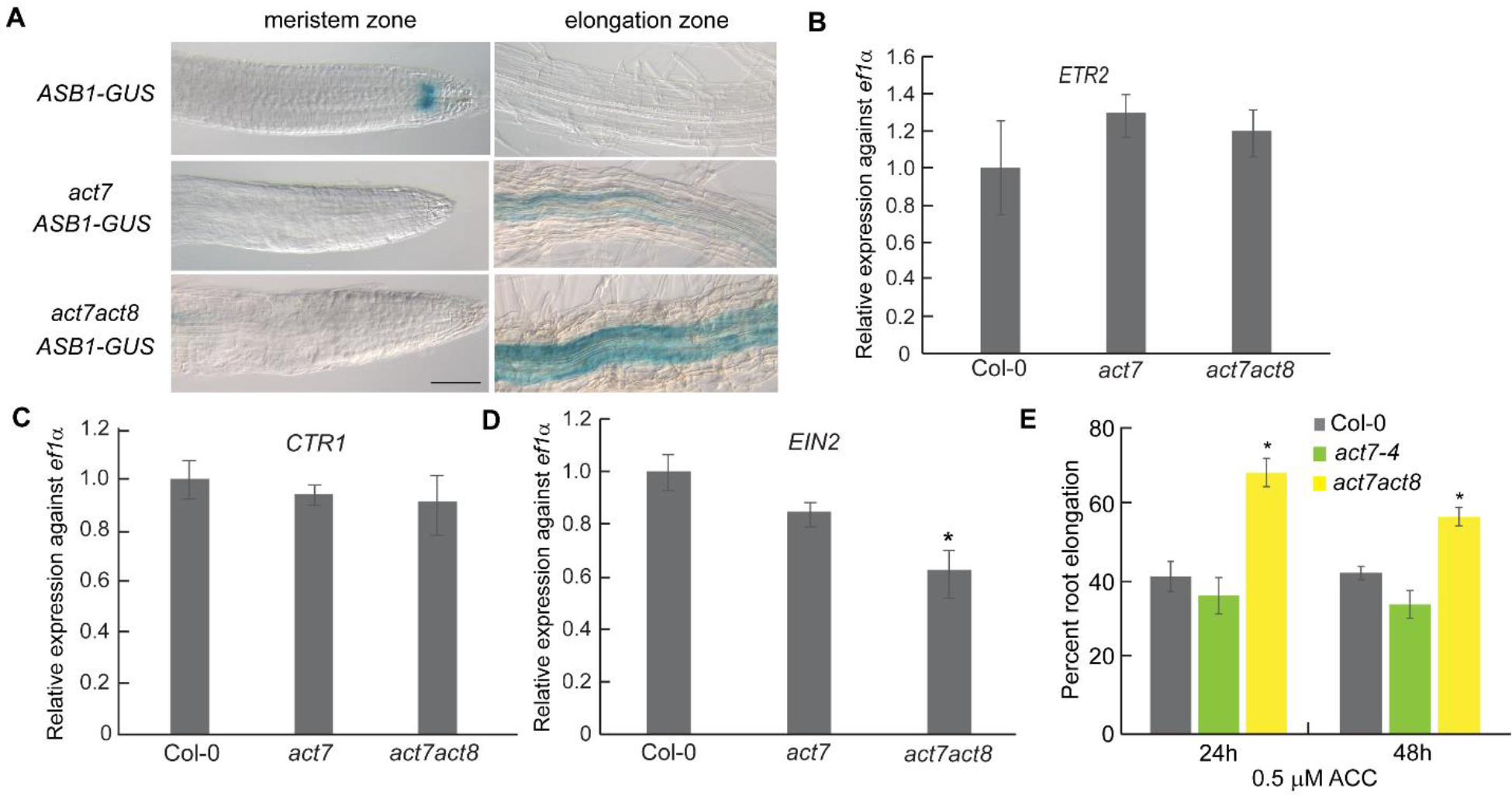
Molecular analysis of ethylene response in *act7* and *act7act8* mutants. (A) Ectopic expression of ethylene-induced auxin biosynthesis gene *ANTHRANILATE SYNTHASE β1* in the mutants. The expression is reduced at the root tip but stimulated in the elongation zone. The images are representative of 20 seedlings from three independent experiments. Bars represent 50 μm. (B-D) Expression analysis of the ethylene signaling genes in *act7* and *act7act8* mutants. qRT-PCR was performed using cDNA prepared from the root samples of seven-day-old seedlings. All the data were normalized against *ef1α*. The data were obtained from three independent biological replicates. (E) Root elongation response assay to ethylene. Seven-day-old light-grown wild-type (Col-0) and mutant seedlings were transferred to new agar plates containing ethylene precursor ACC and grown at 23°C for 48 h. Vertical bars represent mean ± SE of the experimental means from at least four independent experiments (n = 4 or more), where experimental means were obtained from 8–10 seedlings per experiment. Asterisk represents the statistical significance between wild-type and mutants as judged by the Student’s *t*-test (*P<0.5).

### Altered meristem development of *act7* and *act7act8* mutants is independent of cytokinin

Auxin-cytokinin interaction is another major regulator of root meristem development (Dello Ioio et al., 2007). Since *act7* and *act7act8* mutants showed a clear defect in meristematic cell production rate and showed altered auxin response, we next investigated whether the cytokinin response is also affected by the loss of actin isovariants. For this, we observed the expression of cytokinin-specific synthetic marker TCS-GFP, and analyzed the cytokinin-specific genes *ARR1*, *ARR12* (Muller and Sheen, 2007) expression in *act7* and *act7act8* mutants. Unlike auxin markers, we did not observe any alteration in the expression of cytokinin responsive marker or genes in *act7* and *act7act8* mutants (Figs.9A, B). Interestingly, the expression of *Aux/IAA* gene *SHY2*/*IAA3* (Tian and Reed, 1999) was found to be drastically reduced in both mutants. To further confirm that cytokinin response is not altered in these mutants, we performed a root growth assay in presence of exogenous cytokinin. Both *act7* and *act7act8* showed a wild-type like response to kinetin for root elongation inhibition (Fig. 9C). Taken together, these results suggest that the altered meristem development in *act7* and *act7act8* mutants are modulated through a pathway that is independent of cytokinin response.

**Figure 9.**
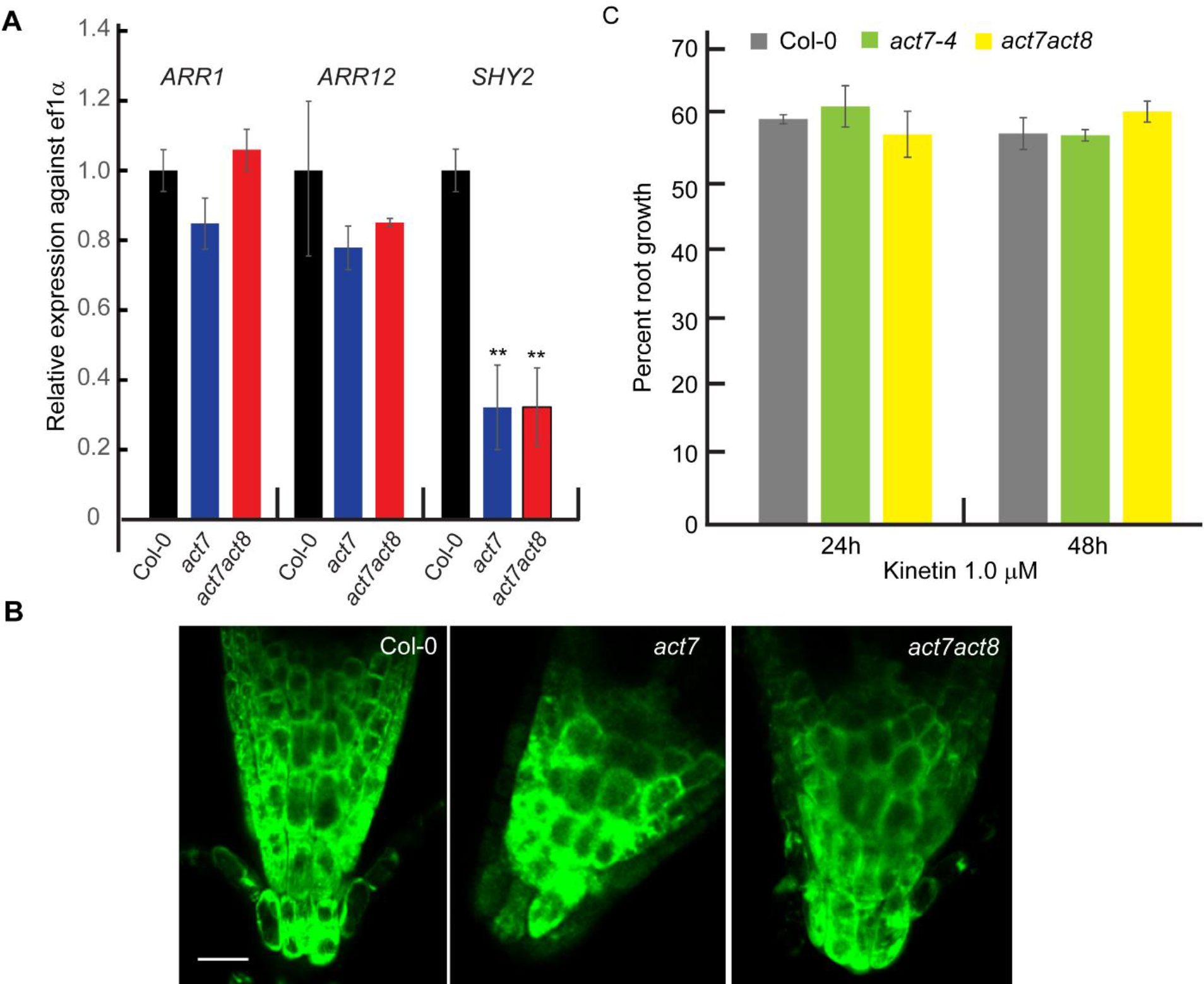
Molecular analysis of cytokinin responsive genes in *act7* and *act7act8* mutants. (A) Expression analysis of the *ARR1*, *ARR12*, and *SHY2/IAA3* in *act7* and *act7act8* mutants. qRT- PCR was performed using cDNA prepared from the root samples of seven-day-old seedlings. All the data were normalized against *ef1α.* The data were obtained from three independent biological replicates. Asterisk represents the statistical significance between wild-type and mutants as judged by the Student’s *t*-test (**P<0.01). (B) Expression of synthetic cytokinin marker *TCS-GFP* is not altered in *act7* and *act7act8* mutants. The images are representative of 20 seedlings from three independent experiments. Bar represents 50 μm. (C) Root elongation response assay to cytokinin. Seven-day-old light-grown wild-type (Col-0) and mutant seedlings were transferred to new agar plates containing Kinetin and grown at 23°C for 48 h. Vertical bars represent mean ± SE of the experimental means from at least three independent experiments (n = 3 or more), where experimental means were obtained from 8–10 seedlings per experiment. No significant difference in root growth was observed between wild- type and mutants as judged by the Student’s *t*-test.

## Discussion

Root organogenesis relies on the primary root meristem development, which in turn is regulated by the coordinated function of cell division, cell differentiation, and cell elongation. Plant hormones auxin, cytokinin, and ethylene have been shown to control this coordination and the local auxin gradient formed by the polar transport of auxin turns out to be the regulatory switch in integrating all these aspects. Earlier, it was shown that cytokinin promotes root cell differentiation by suppressing the auxin distribution in the meristem (Dello Ioio et al., 2008a; Růzǐčka et al., 2009), while the formation of local auxin gradient at the root apex promotes the cell division (Brumos et al., 2018b). Ethylene also affects root development through modulating auxin transport and auxin gradient (Růzicka et al., 2007; Lewis et al., 2011). The formation of auxin gradient depends on the polar transport of auxin, which is regulated by the intracellular trafficking mediated correct localization of the cellular auxin efflux carriers PINs (Luschnig et al., 1998; Geldner et al., 2001). The cell cytoskeleton component actin plays a major role in facilitating the trafficking of auxin efflux carriers by providing tracks for the movement of these proteins (Geldner et al., 2001; Muday and Murphy, 2002; Rahman et al., 2007; Kleine-Vehn et al., 2008). However, what remains obscure is which type of actin contributes to this trafficking process and whether they directly regulate the cellular auxin gradient formation and thereby root meristem development. Taking advantage of the vegetative class actin isovariant mutants, in this study we answered these questions. Our results demonstrate that root meristem maintenance is largely dependent on the ACT7 class actin filament (Fig. 2). Loss of ACT7 results in fragmented and depolymerized actin that affects the expression and trafficking of a subset of auxin efflux carriers mainly PIN1, PIN2, and PIN7, and subsequently the polar transport mediated auxin gradient at the root tip (Figs. 1, 3, 4, 5, 6). The consequence of this cascade results in reduced cell division and shorter root meristem (Fig. 2). These findings are consistent with the previous results showing that disruption of actin, by actin depolymerization drug Latranculin B resulted in reduced cell division and inhibition of primary root elongation (Baluška et al., 2001; Rahman et al., 2007), and *act7* mutant shows a dwarf seedling phenotype including a severe reduction in the primary root elongation (Gilliland et al., 2003). The actin structure, trafficking, expression of PIN proteins, and root meristem development were largely unaffected in absence of other vegetative class actin isovariants ACT2 or ACT8. Collectively, these results suggest that the intact actin cytoskeleton maintained by ACT7 isovariant plays an essential role in root meristem development.

Although the subclass-specific Arabidopsis actin isovariants share high homology, there are 28-29 amino acid substitutions between *At*ACT7 and *At*ACT2 and *At*ACT8 (Mcdowell et al., 1996). The differential activities of ACT7 and ACT2/8 in root development may be attributed to the subclass variance and subtle changes in biochemical properties. For instance, ACT2/8 clade groups with genes from maize and rice compared to ACT7, while ACT7 is closer to the genes from pea, carrot, potato, and pine, suggesting that they originated from different lineages. The differences between these two subclasses are also evident in their amino acid substitutions, which have the potential to alter actin-actin or actin-actin binding protein interactions (McDowell et al., 1996). Differential biochemical properties of ACT7 and ACT2 have been reported in terms of constructing unique filament arrays depending on the cell type or tissues, and binding to specific actin binding proteins (Kijima et al., 2018). It has also been reported that actin binding proteins, profilin 1 and profilin 2 inhibit the polymerization of ACT7, while the effect is minimal on ACT2 (Kijima et al., 2016). Future research focusing on identifying the isovariant specific binding proteins will further enhance our understanding of the underlying mechanisms.

The analysis of the actin isovariant mutants supported the previous claims that the trafficking of the auxin efflux carriers PIN1 and PIN2 is actin-dependent (Geldner et al., 2001; Muday and Murphy, 2002; Rahman et al., 2007). It further clarifies that ACT7 isovariant is the main actin modulating this process as PIN1 and PIN2 trafficking as well as the auxin gradient at the tip is not affected at all by the loss of ACT2 and ACT8 (Figs. 3, 4, 6). Interestingly, PIN4, PIN7, and AUX1 trafficking are not affected by the loss of ACT7, suggesting that not all auxin carrier proteins trafficking or membrane localization are actin-dependent (Fig. 5). The ACT7 mediated PIN1 and PIN2 trafficking is possibly regulated by the ABCB chaperone TWISTED DWARF1 (TWD1) as ACT7 was found to be an indirect interactor of TWD1 (Zhu et al., 2016). However, there could be other ACT7 interacting proteins that may regulate this process and need to be identified. The finding that only one actin isovariant ACT7 affects the root meristem developmental process through modulating auxin distribution is not inconsistent as it was previously shown that ACT7 affects the primary root growth and response to auxin during callus formation (Kandasamy et al., 2001; Kandasamy et al., 2009).

The reduced meristem growth in *act7* was found to be linked to the inhibition of cell division and cell elongation (Fig. 2, Table 1). Our results suggest that this growth reduction and the inhibition of cell division are linked to the auxin maximum at the root tip. In *act7* mutant, we observed a depletion in auxin maximum, while in *act7act8* mutant the auxin maximum was absent at the root tip (Fig. 6). The reduced auxin maxima in *act7* and *act7act8* mutants are a consequence of both reductions in PIN1 and PIN2 regulated polar transport of auxin, and the auxin synthesis at the root tip (Figs. 7 and 8). This difference in auxin maxima is reflected in their root meristem development and cell division where *act7act8* shows a more severe phenotype compared with *act7* (Fig. 2; Table 1). Interestingly, auxin efflux double mutant *pin1pin2* showed a similar meristem developmental defect observed in *act7* or *act7ac8* confirming that PIN transporters-mediated auxin transport plays an important role in this process (Blilou et al., 2005). Although *act7* and *act7act8* mutants show less or no auxin maxima at the root tip, they showed auxin response in the elongation zone confirming that they retain the capability to respond to auxin. Consistently, exogenous application of auxin results in an increased response in the elongation zone (Fig. 6). These results are consistent with the idea that local auxin maximum at root tip is an absolute requirement for meristematic cell division and meristem growth (Grieneisen et al., 2007; Brumos et al., 2018b).

Myosin XI is an actin-based motor protein, and this dynamic actin-myosin XI system plays a major role in subcellular organelle transport and cytoplasmic streaming (Duan and Tominaga, 2018). Interestingly, earlier it was demonstrated that Myosin XI-K orchestrates root organogenesis through affecting polar auxin transport, auxin response, and cell division (Abu-Abied et al., 2018). The similar mode of action of ACT7 and Myosin XI in modulating root organogenesis raises a possibility that loss of ACT7 may alter the myosin XI activity, which in turn affects the cell division capability of the root. Proof of this intriguing possibility awaits further experimentation.

Loss of *act7* and *act8* also affected the root ethylene response. Ethylene-specific auxin biosynthesis gene *ASβ1* was found to be downregulated in both *act7* and *act7act8* mutants (Fig. 8). On the other hand, expressions of the genes that regulate the ethylene response pathway were largely unaltered in these mutants except *EIN2*, which was downregulated in *act7act8* mutant background. Consistently, *act7act8* mutant showed resistance to ACC-induced root growth inhibition (Fig. 8). These results suggest that the root ethylene response depends on the extent of the auxin gradient present at the root tip. This idea is consistent with the observation that although reduced compared to wild-type, *act7* still retains transport-mediated auxin gradient at the root tip, which is sufficient for ethylene response (Figs. 6 and 8). In contrast, in *act7act8*, the auxin gradient at the root tip is completely depleted and *act7act8* shows reduced ethylene response. Collectively, these results suggest that the altered root meristem development in *act7* and *act7act8* mutants are consequences of loss of both auxin and ethylene responses.

Involvement of auxin-cytokinin interaction in root meristem development has been well studied and shown to be controlled through a balance between cell division and cell differentiation mediated by a simple regulatory circuit converging on Aux/IAA family protein SHY2/IAA3 (Dello Ioio et al., 2008a; Moubayidin et al., 2010b). The primary cytokinin-response transcription factors *ARR1* and *ARR12* activate the *SHY2/IAA3*, a repressor of auxin signaling, that negatively regulates the expression of *PIN1*, *PIN3,* and *PIN7* genes and thereby alters the auxin distribution in the root meristem (Dello Ioio et al., 2008a; Růzǐčka et al., 2009; Moubayidin et al., 2010b). To understand whether the altered auxin maximum at the root tip and the altered root meristem development of *act7 a*nd *act8* is a consequence of altered cytokinin response, we investigated the expression of *ARR1*, *ARR12,* and *SHY2/IAA3* in the mutant background. Neither *ARR1* nor *ARR12* expression was altered in *act7* or *act7act8* double mutant. Consistently, we also did not observe any change in the synthetic cytokinin marker *TCS-GFP* (Fig. 9). Interestingly, we found that *SHY2/IAA3*, which belongs to the primary auxin-responsive gene family *Aux/IAA* (Abel and Theologis, 1996) was downregulated in both *act7* and *act7act8* mutants (Fig. 9). *IAA3* has also been shown to directly influence the auxin-regulated Arabidopsis root development (Tian and Reed, 1999). These observations confirm that the altered root meristem development observed in loss of ACT7 mutant is linked to altered auxin response. *ARR1*, *ARR12* regulated *SHY2*/*IAA3* pathway is more dominant in older seedlings compared with younger seedlings as at the early seedling stage only *ARR12* is expressed and *SHY2* level is relatively low (Moubayidin et al., 2010b). Consistently, auxin-regulated cell division predominates over cytokinin-regulated cell differentiation to shape the meristem development at the early seedling stage (Chapman and Estelle, 2009; Moubayidin et al., 2010b). The observation that the altered root meristem development in the loss of ACT7 is primarily dependent on auxin response and independent of cytokinin is consistent with the above findings as we characterized the root meristem at the early stage of development.

In conclusion, in the present study, we demonstrated that primary root meristem development is controlled by actin isovariant ACT7 mediated auxin redistribution in the root tip and primarily modulated by auxin-ethylene responses but independent of cytokinin.

## Materials and methods

### Plant materials and growth conditions

All lines are in the Columbia background of *Arabidopsis thaliana* (L.). *ASB1-GUS* was a gift of Jose Alonso (University of North Carolina, Raleigh, USA, Stepanova et al., 2005), PIN2-GFP (Xu and Scheres, 2005) was a gift from B. Scherers (Wageningen University, The Netherlands), GFP-ABD2–GFP (Wang et al., 2007) was a gift from E.B. Blancaflor (Samuel Roberts Noble Foundation, Ardmore, OK, USA). *IAA2:GUS* and AUX1-YFP were gifts of Malcolm Bennett (Swarup et al., 2007). PIN4-GFP and PIN7-GFP were gifts of Gloria Muday (Wake Forest University, NC, USA, (Vieten et al., 2005; Laskowski et al., 2008). Columbia-0, PIN1-GFP, and *aux1-7* were obtained from the Arabidopsis Biological Resource Center (Columbus, OH, USA). The *act8-2, act7-4* a*ct7act8, act2-1*, and *act2act8* were gifts of R. Meagher (University of Georgia, Athens, Georgia, Kandasamy et al., 2009). Various marker lines in actin isovariant mutant background were generated in the laboratory by crossing, and independent homozygous lines for the mutation and expressing the GFP or GUS reporter were identified by screening for fluorescence, GUS assay, and seedling phenotype.

Seeds were sterilized in 20% kitchen bleach (Coop Clean Co., http://www.coopclean.co.jp) for 15 minutes and washed 3 times in sterilized dH2O. Surface-sterilized seeds were placed in round, 9 cm Petri plates on modified Hoagland’ s medium (Baskin and Wilson, 1997) containing 1% w/v sucrose and 1% w/v agar (Difco Bacto ager, BD laboratories; http://www.bd.com). Two days after stratification at 4°C in the dark, plates were transferred to the growth chamber (NK system, LH- 70CCFL-CT, Japan, http://www.nksystems.co.jp/) at 23°C under continuous white light at an irradiance of about 100 μmol m^-2^ s^-1^. The seedlings were grown vertically for 5 or 7 days.

### Chemicals

IAA was purchased from Sigma-Aldrich Chemical Company (http://www.sigmaaldrich.com). Other chemicals were purchased from Wako Pure Chemical Industries (http://www.wako-chem.co.jp/). FM4-64 was purchased from Invitrogen (http://www.thermofisher.com). [^3^H] IAA (20 Ci mmol^−1^) was purchased from American Radiolabeled Chemicals (http://www.arcinusa.com).

### Growth, cell length and cell production rate assays

Root elongation rate was measured by scoring the position of the root tip on the back of the Petri plate once per day. Cortical cell length was measured using a light microscope (Diaphot, Nikon, www.nikon.co.jp) equipped with a digital camera control unit (Digital Sight [DS-L2], Nikon) as described earlier (Rahman et al., 2007). To ensure newly matured cells were scored, no cell was measured closer to the tip than the position where root hair length was roughly half - maximal. The length of twenty mature cortical cells was measured from each root, with eight roots used per treatment. The cell production rate (cells day^-1^) was calculated by taking the ratio of root elongation rate (mm day^-1^) and average cell length (μm) for each individual and averaging over all the roots in the treatment. The results were obtained from at least three biological replicates per genotype.

### GUS staining, immunostaining, and live-cell imaging

GUS staining was performed as described earlier (Okamoto et al., 2008). In brief, 5-d-old vertically grown seedlings were used for GUS assay. Seedlings were transferred to GUS staining buffer (100 mM sodium phosphate pH 7.0, 10 mM EDTA, 0.5 mM potassium ferricyanide, 0.5 mM potassium ferrocyanide, and 0.1 % Triton X-100) containing 1mM X-gluc and incubated at 37°C in the dark for various time length as mentioned in the figure legends. To observe the effect of exogenous auxin on GUS expression, the seedlings were incubated in 1 μM IAA for 3h and subjected to GUS staining as described above. Roots were cleared as described earlier (Shibasaki et al., 2009). The roots were imaged with a light microscope (Nikon Diaphot) equipped with a digital camera control unit (Digital sight, DS-L2; Nikon).

Actin immunostaining was performed using the protocol described earlier by (Bannigan et al., 2006; Rahman et al., 2007 ) with modification. In brief, Five-day-old Arabidopsis seedlings were fixed in PIPES buffer (50 mM PIPES, 4% paraformaldehyde, 1.2% glutaraldehyde, 1 mM CaCl2, 0.4 mM maleimidobenzoyl-N-hydroxy succinimide) for 60 min, followed by rinsing in PME buffer (50 mM PIPES, 5 mM EGTA, and 2 mM MgSO4) 3 times for 10 min each. After the PME washing, the seedlings were extracted for 60 min in PME buffer containing 1% Triton X-100. The seedlings were then subjected to digestion for 15 min in PBS with 0.001% pectolyase and 0.01% pectinase, and rinsed three times for 5 min in PME with 10% glycerol and 0.002% Triton X-100. Seedlings were permeabilized by incubating them at −20°C in methanol for 30 min, followed by rehydration in PBS for three times 5 min each. Seedlings were incubated in the primary mouse monoclonal anti-(chicken gizzard) actin, clone 4 (Millipore, http://www.millipore.com) diluted 1:200 in PBS, 1% BSA, and 0.01% sodium azide (PBA) overnight. After the incubation, the seedlings were washed for three times 5 min each and then incubated in the secondary antibody Cy-3-goat anti-mouse IgG (1:200; Jackson Immunoresearch, http://www.jacksonimmuno.com). The imaging was performed on a Nikon laser scanning microscope (Eclipse Ti equipped with Nikon C2 Si laser scanning unit, http://www.nikon.com) equipped with a ×60 oil immersion objective.

To image GFP, YFP and FM 4-64-stained roots, five-day-old seedlings were mounted in liquid growth medium on a cover glass for observation on a Nikon laser scanning microscope (Eclipse Ti equipped with Nikon C2 Si laser scanning unit) and imaged with either a ×20 dry or ×40 water immersion objectives. The images were taken using the same confocal settings for each set of experiments. All the experiments were repeated at least 3 times.

### Actin quantification

Five-day-old seedlings of ABD2-GFP and ABD2-GFP in different actin mutant backgrounds were subjected to imaging with the Nikon laser scanning microscope (Eclipse Ti equipped with Nikon C2 Si laser scanning unit) using a 40x water immersion objective with a numerical aperture of 1.25, and the same laser set up. All the experiments were repeated at least three or four times. Actin quantification was performed using ImageJ software (http://rsb.info.nih.gov/ij/) as described earlier (Higaki et al.,2010, Takashi et al., 2014, Parveen and Rahman, 2021). Before measurements of actin parameters, the confocal images were skeletonized using the ImageJ plug-in LPX LineExtract (Ueda et al., 2010). Using the processed images, we measured four different parameters: occupancy for density, skewness for bundling, Δθ for orientation, and NormAvgRad for parallelness.

### Root meristem length measurement

For root meristem imaging, seven-day-old seedlings were stained in 2μM FM 4-64 for 15 minutes and subjected to imaging under Nikon laser scanning microscope (Eclipse Ti equipped with Nikon C2 Si laser scanning unit) using the same settings. Images and data are representative of at least 3-4 biological replicates. In each run, 6-8 seedlings were imaged.

### Auxin transport assay

Five-day-old vertically grown *Arabidopsis* seedlings were used for auxin transport assay. Auxin transport was measured as described earlier (Shibasaki et al., 2009). In brief, a donor drop was prepared by mixing 0.5 μM [^3^H] IAA (3.7 MBq ml^−1^) in 1.5% agar containing MES buffer solution. The donor drop was placed on the edge of the root tip for the shootward auxin transport assay or at the root-shoot junction for the rootward transport assay. Plates were then incubated vertically at 23°C for 2h for shootward auxin transport assay and 6h for rootward auxin transport assay. For measurement of auxin transport, 5-mm root segments away from the apical 2 mm root tip and 2 mm away from the root-shoot junction were carefully cut and soaked overnight in 4 ml of liquid scintillation fluid (Ultima Gold, PerkinElmer, USA), and the radioactivity was measured with a scintillation counter (model LS6500, Beckman ACSII; USA Instruments, Fullerton, CA). Data were obtained from at least three biological replicates.

### Gene expression analysis

RNA was extracted from 5-day-old vertically grown *Arabidopsis thaliana* seedlings root tissue using RNeasy Mini Kit (Qiagen, www.qiagen.com) with on-column DNA digestion to remove residual genomic DNA using RNase-free DNase according to manufacturer’s protocol. Extracted RNA was tested for quality and quantity. Each RNA concentration was normalized with RNase-free water. 500 ng RNA was used to synthesize cDNA using Rever Tra Ace qPCR RT master mix (Toyobo, Japan, www.toyobo-global.com). Quantitative PCR reactions were performed using the Takara TP-850 thermal cycler (Takara Bio, Japan, www.takara-bio.com) and THUNDERBIRDTM SYBR® qPCR Mix from Toyobo (https://www.toyobo.co.jp). The reaction was performed as per manufacturer’s instruction. For quantification of gene expression, we used the 2^-ΔΔ^CT (cycle threshold) method (Livak and Schmittgen, 2001) with normalization to the *ef1α* expression. Data were obtained from three biological replicates. Primers used for the gene expression analysis are listed in Supplemental Table 1.

### Statistical analysis

Results are expressed as the means ± SE from the appropriate number of experiments as mentioned in the figure legends. Statistical significance was analyzed using a two-tailed Student’s *t*-test for comparison between wild type and mutants, and analysis of variance (ANOVA) followed by the Tukey-Kramer test for comparisons between multiple genotypes.

## Acknowledgments

The authors thank Rich Meagher ( University of Georgia, Athens, USA), Jose Alonso (North Carolina State University, Raleigh, USA), B. Scheres (Wageningen University, The Netherlands), E.B. Blancaflor (Samuel Roberts Noble Foundation, Ardmore, OK, USA), Malcolm Bennett (University of Nottingham, UK), and Gloria Muday (Wake Forest University, NC, USA). This work was funded by JSPS Kakenhi (Grant Number 23012003 A.R.). We thank Michiko Saito for the technical assistance.

## Author Contribution

A.R., A.A.R. designed the experiments. T.N. and K.S. performed the experiments. A.R. supervised the experiments. All the authors analyzed the data. A.R. and A.A.R. wrote the manuscript.

