## Supplemental Information for "Actin isovariant ACT7 controls root meristem development in Arabidopsis through modulating auxin and ethylene responses"

**Supplemental Table 1**: Primers used for gene expression analysis in this study.

| | |
|---|---|
| PIN1-F | ATCTTCACATGTTTGTGTGG |
| PIN1-R | TCGTCTTTGTTACCGAAACT |
| PIN2-F | AGATGCCAACGATAATGAGT |
| PIN2-R | AGTAATCACCTGAACGATGG |
| PIN3-F | AGATCTGACCAAGGTGCTAA |
| PIN3-R | CCTAGACCTGTCTTGGATTG |
| PIN4-F | ACTTCAACCCAAAATCATTG |
| PIN4-R | GTGGGATGCACATTGTACT |
| PIN7-F | AGTTGATAATGGAGCCAATG |
| PIN7-R | TTATGAGTTTCCTCCACACC |
| AUX1-F | CAGATCAGGTAAACGGAAAC |
| AUX1-R | TCCAGCTTCCTAGTAAACCA |
| ARR1-F | GTCAAGACACAACACGACAG |
| ARR1-R | TGTATCCGTAGCCACTCTCT |
| ARR12-F | CAGCTTCAGACAAACAACAA |
| ARR12-R | GTTGCGTAGAGAAGCTAGGA |
| SHY2-F | GGGTCAAGGAATCTATGTGA |
| SHY2-R | CCCACAGAGAATTTGAACAT |
| ETR2-F | AGAGAAACTCGGGTGCGATGT |
| ETR2-R | TCACTGTCGTCGCCACAATC |
| CTR1-F | GCTGCATTACCACAAAAGAGG |
| CTR1-R | AGTAAAGCTCGATGTCTGCA |
| EBF2-F | TGATGTTGGTCTTGGTGCTGTTGC |
| EBF2-R | ATTCCAGGACACCGTGAAAGGTCA |
| EIN2-F | GGAGGGTATGGTGCGTCTTA |
| EIN2-R | TGTGGCAAACTGTAGGCATC |

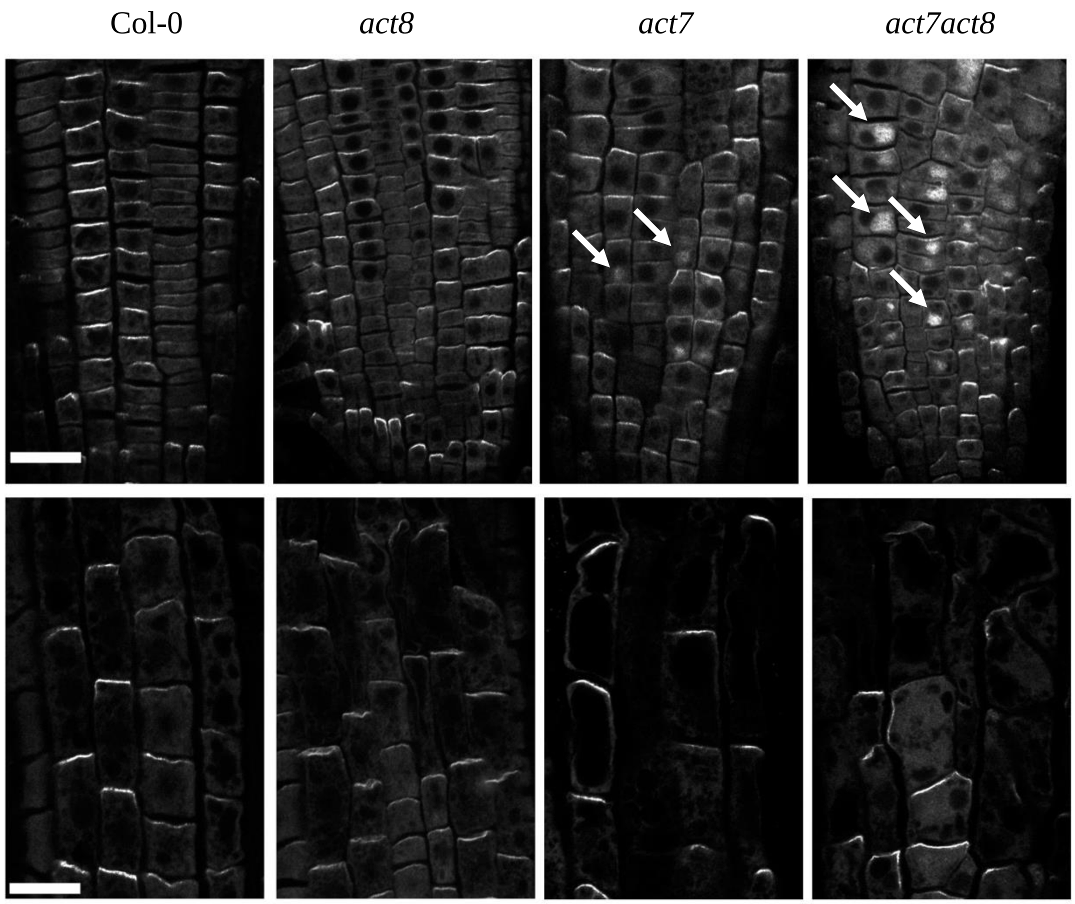


**Supplemental Figure 1**: Localization of PIN2 in actin mutants. Five-day-old seedlings were fixed and processed for immunofluorescence with an anti-PIN1 antibody. The upper panel represents the meristem zone and the lower panel represents transition zone, respectively. Five-day-old seedlings were fixed and processed for immunofluorescence with an anti-PIN1 antibody. The images are single stack and representative of three biological replicates. Imaging was performed with same confocal settings. Arrowheads indicate the intracellular agglomeration of PIN2. Bar=50 μm

**Method for PIN immunostaining:**

PIN localization was performed using the method described earlier (Rahman et al., 2007). Five-day-old seedlings were used for immunostaining. The primary antibody was anti-PIN2 (1:100 dilution) generated in our lab. The secondary antibody was Cy-3 goat anti-rabbit IgG (1:200, Jackson Immunoresearch, **http://www.jacksonimmuno.com/**). All imaging was done on a confocal laser microscope, (Nikon laser scanning microscope, Eclipse Ti equipped with Nikon C2 Si laser scanning unit, www.nikon.co.jp) and imaged with 60× oil-immersion objective. The images were taken using the same confocal settings for each set of experiments. All the experiments were repeated at least 3 times.
